# Genetic variants related to antihypertensive targets inform drug efficacy and side effects

**DOI:** 10.1101/460543

**Authors:** Dipender Gill, Marios K. Georgakis, Fotios Koskeridis, Lan Jiang, Qiping Feng, Wei-Qi Wei, Evropi Theodoratou, Paul Elliott, Joshua C. Denny, Rainer Malik, Evangelos Evangelou, Abbas Dehghan, Martin Dichgans, Ioanna Tzoulaki

## Abstract

**Background:** Drug effects can be investigated through natural variation in the genes for their protein targets. We aimed to use this approach to explore the potential side effects and repurposing potential of antihypertensive drugs, which are amongst the most commonly used medications worldwide.

**Methods:** We identified genetic instruments for antihypertensive drug classes as variants in the gene for the corresponding target that associated with systolic blood pressure at genome-wide significance. To validate the instruments, we compared Mendelian randomisation (MR) estimates for drug effects on coronary heart disease (CHD) and stroke risk to randomised controlled trial (RCT) results. Phenome-wide association study (PheWAS) in the UK Biobank was performed to identify potential side effects and repurposing opportunities, with findings investigated in the Vanderbilt University Biobank (BioVU) and in observational analysis of the UK Biobank.

**Findings:** We identified suitable genetic instruments for beta-blockers (BBs) and calcium channel blockers (CCBs). MR estimates for their effect on CHD and stroke risk respectively were comparable to results from RCTs against placebo. PheWAS in the UK Biobank identified an association of the CCB genetic risk score (scaled to drug effect) with increased risk of diverticulosis (odds ratio [OR] 1.23, 95%CI 1.10-1.38), with a consistent estimate found in BioVU (OR 1.16, 95%CI 0.94-1.44). Association with diverticulosis was further supported in observational analysis of CCB use in the UK Biobank (OR 1.08, 95%CI 1.02-1.15).

**Interpretation:** We identified valid genetic instruments for BBs and CCBs. Using genetic and observational approaches, we highlighted a previously unreported potential detrimental effect of CCBs on risk of diverticulosis. This work serves as a proof of concept that investigation of genetic variants can offer a complementary approach to exploring the efficacy and side effects of anti-hypertensive medications.

**Funding:** Wellcome Trust.

## Research in context

### Evidence before this study

Hypertension is a leading cause of mortality worldwide and use of antihypertensive drugs is widespread. Drug effects can be investigated by studying genetic variants in the genes of their corresponding protein targets. The availability of large-scale genome-wide association study meta-analyses for blood pressure has now made this possible for antihypertensive agents.

### Added value of this study

This study searched for genetic variants to instrument the effect of first-line antihypertensive drugs. Mendelian randomization showed that instruments identified for the beta-blocker (BB) and calcium channel blocker (CCB) classes produced estimates for risk of coronary heart disease and stroke comparable to those obtained in clinical trials. Phenome-wide association study in the UK Biobank corroborated the established efficacy of these agents in preventing diseases related to hypertension, and additionally highlighted a detrimental association of the CCB genetic risk score with diverticulosis. A consistent estimate for this apparent detrimental association was found in the Vanderbilt University Biobank and was further supported in observational analysis of the UK Biobank.

### Implications of all the available evidence

The use of genetic variants offers a complementary approach to explore the efficacy, side effects and repurposing potential of the BB and CCB classes of anti-hypertensive medication. Using genetic and observational approaches, we find evidence of a detrimental effect of CCBs on risk of diverticulosis. Given the widespread use of this drug class, these findings could have notable clinical implications and warrant further investigation.

## Introduction

In 2015, the 874 million adults worldwide estimated to have a systolic blood pressure (SBP) of 140mmHg or higher accounted for 106 deaths per 100,000 and loss of 143 million disability-adjusted life years (1), making hypertension a leading cause of mortality and morbidity. Blood pressure lowering through lifestyle modification or pharmacological treatment can significantly decrease cardiovascular risk, with every 10mmHg reduction estimated to decrease risk of all-cause mortality by 13% (2).

The pharmacological treatment of hypertension is founded on strong evidence, underpinned by a large number of outcome-based randomised controlled trials (RCTs) that have identified several drug classes to be effective for lowering blood pressure (3). However, RCTs based on clinical outcomes have limitations (4); they are largely restricted to older or high-risk patients and have a relatively short duration of follow-up, rarely beyond five years (5). Therefore, recommendations for treatment are often based on extrapolation of the available evidence, with known side effects frequently limited to relatively common outcomes captured in RCTs (6). At the same time, particular drug treatments for hypertension may have beneficial effects beyond their blood pressure lowering properties (6), thus offering potential for repurposing. However, observational research used to study such opportunities suffers from well-characterised biases, including confounding by indication (7).

With the growing availability of genome-wide association study (GWAS) meta-analyses, it is becoming increasingly feasible to study drug effects by investigating genetic variants in the genes of their protein targets, as has previously been applied to lipid lowering drugs (7). In a proof of concept study, we searched for human genetic variation within genes corresponding to first-line pharmacological agents for hypertension to proxy the effects of these treatments. We investigated the validity of this approach by exploring consistency in Mendelian Randomisation (MR) estimates for their effect on coronary heart disease (CHD) and stroke risk with corresponding RCT findings. To offer insight towards their adverse effect profiles or repurposing potential, we undertook phenome-wide association study (PheWAS) analyses with replication in an external dataset, and further investigated our findings in observational analysis of the UK Biobank.

## Methods

### Genetic instrument selection

First-line antihypertensive drugs for study were selected based on recent consensus guidelines (6): angiotensin-converting-enzyme inhibitor (ACEI), angiotensin receptor blocker (ARB), beta-blocker (BB), calcium channel blocker (CCB) and thiazide diuretic (TD). Genes encoding the targets of these drugs were identified using the DrugBank database (8), with promoter and enhancer regions identified using the GeneHancer database in the GeneCards online platform (9). Genetic association estimates for SBP were obtained from a GWAS meta-analysis of 757,601 individuals with European ancestry drawn from the UK Biobank and the International Consortium of Blood Pressure GWAS meta-analysis (10), where correction was made for antihypertensive medication use by adding 15mmHg to the SBP of participants receiving medication, with further adjustment for body mass index (BMI) (10). In sensitivity analyses, we also used a GWAS of SBP on approximately 337,000 White British individuals in the UK Biobank, without correction for medication use or adjustment for BMI (11). Instruments for the genetic effect of lower SBP through antihypertensive drug targets were selected as single-nucleotide polymorphisms (SNPs) in corresponding genes, promoter regions or enhancers, that were associated with SBP at genome-wide significance (*P*<5 x 10^−8^) and clumped to a linkage disequilibrium (LD) threshold of r^2^<0.1 using the 1000G European reference panel. The R^2^ and F statistics were used to estimate the variance in SBP explained and strength of each SNP respectively (12).

### Mendelian randomisation

Only antihypertensive drugs where 3 or more SNPs were identified as instruments using the larger SBP GWAS were taken forward to MR analysis, in order to allow us to investigate for potential violations of the requisite MR assumptions (13). Genetic association estimates for CHD were obtained from the CARDIoGRAMplusC4D Consortium’s 1000 Genomes-based trans-ethnic meta-analysis of 60,801 cases and 123,504 controls (14). Estimates for stroke risk were obtained from the MEGASTROKE Consortium’s trans-ethnic meta-analysis of 67,162 cases of any stroke and 454,450 controls (15). Ethical approval was obtained in each of the participating studies contributing to these GWAS meta-analyses and only published summary data were used, thus further ethical approval was not required.

In the main MR analysis, estimates for each SNP were derived using the Wald ratio, with standard errors estimated using second order weights to allow for measurement error in both the exposure and outcome estimates (16). Overall MR estimates were calculated by pooling individual MR estimates for each SNP instrumenting a specific drug target using fixed-effects inverse-variance weighted (IVW) meta-analysis (16), and were scaled to the estimated effect of the corresponding drug target on SBP in RCTs: 21.14mmHg decrease for ACEI, 9.51mmHg decrease for BB, 8.90mmHg decrease for CCB, and 12.56mmHg decrease for (low-dose) TD (3). After conversion to relative risk (RR) estimates with the baseline incidences of CHD and stroke taken as 0.042 and 0.041 respectively (2), MR results were compared with estimates from a recent Cochrane systematic review and meta-analysis of RCTs that investigated the effect of first-line antihypertensive drugs against placebo (3).

### Investigation of pleiotropy

MR estimates may be biased when the genetic variants used as instruments affect the outcome through a pleiotropic pathway that is independent of the exposure. The PhenoScanner database of SNP-phenotype associations was used to explore whether any of the selected instrument SNPs or proxies with LD r^2^<0.8 (using a 1000G reference panel) were also associated at genome-wide significance (*P*<5×10^−8^) with traits that may potentially be exerting pleiotropy (17), and any such SNPs were excluded in sensitivity analyses.

We also incorporated statistical evaluation of possible pleiotropy (13). Heterogeneity in the MR estimates generated by different instrument SNPs can be used to indicate such pleiotropy (13), which we identified through a significant Cochran’s Q test (*P*<0.05) or an I^2^ measure of heterogeneity >30%. We also performed MR statistical sensitivity analyses that are more robust to the inclusion of pleiotropic variants. Firstly, we used the weighted median estimator, which obtains an overall MR estimate by ordering individual SNP MR estimates by their magnitude weighted for their precision, and is reliable when more than half the information for the analysis comes from valid instruments (18). Secondly, we used the MR-Egger technique, which regresses the SNP-outcome estimates against the SNP-exposure estimates, weighted for the precision of the SNP-outcome estimates to give a reliable MR estimate and test for the presence of directional pleiotropy in scenarios where any pleiotropic effect of the instruments is independent of their association with the exposure (19).

Finally, we conducted MR-PRESSO, which performs a zero-intercept regression of the SNP-outcome estimates against the SNP-exposure estimates to test, using residual errors, whether there are outlier SNPs (*P*<0.05), and whether removing these changes the MR estimates generated (20). MR-PRESSO generally requires that at least half of the instruments used do not relate to the outcome independently of the exposure (20). Statistical sensitivity analyses in MR suffer from low power (13), and as such no formal statistical significance threshold was set for these.

### Phenome-wide association study (PheWAS)

The UK Biobank served as the cohort for the PheWAS. It is a prospective study comprising 503,317 individuals aged 40-69 recruited 2006-2010 (21). The participants provided a wide range of self-reported information, with blood samples collected for biochemical tests and genotyping, and a range of physical measurements performed as previously described (21). Individuals were linked retrospectively and prospectively to Hospital Episode Statistics (HES).

PheWAS was restricted to participants of self-reported European descent, with random exclusion of one participant from each pair of relatives based on a kinship coefficient >0.0884. For antihypertensive drugs where 3 or more SNPs were identified as instruments, we used PLINK to construct a genetic risk score (GRS) for each individual, weighted for the effect of each comprising SNP on SBP (22). The GRSs were scaled to the estimated effect of the corresponding drug target on SBP in RCTs (3). The 9^th^ and 10^th^ revisions of the International Classification of Diseases (ICD) were used to define cases based on inpatient HES data. The phecode grouping system was used to align diagnoses used in clinical practice with genomic analysis (23). A series of case-control groups were generated for each phecode, with controls identified as individuals with no record of the respective outcome and its related phecodes (23). Analysis was performed using logistic regression after adjusting for age, sex and first four genetic principal components. We only considered outcomes that had a minimum of 200 cases in order to maintain sufficient statistical power to identify associations with common variants (24). A 5% threshold with the false discovery rate (FDR) method was used in ascertaining the statistical significance of associations, to correct for multiple testing of correlated phenotypes. As for the MR analysis, we also performed sensitivity analyses using genetic instruments derived from the SBP GWAS that did not correct for medication use or adjust for BMI, and after excluding any SNPs with potentially pleiotropic associations at genome-wide significance that were identified using PhenoScanner (17).

We further investigated any FDR significant PheWAS associations for non-cardiovascular conditions in the Vanderbilt University Biobank (BioVU), for which genetic data on approximately 55,000 individuals are linked to a de-identified Electronic Health Record system (25). Similarly to the main PheWAS, a weighted GRS was constructed and logistic regression with the outcome was performed after adjusting for age, sex and first three principal components. The analysis was restricted to individuals identified as White, with controls based on the same exclusions as the main PheWAS. Results between UK Biobank and BioVU analysis were pooled using a fixed-effects meta-analysis model.

### Observational analysis

We further explored significant PheWAS associations for non-cardiovascular conditions in observational analysis of the UK Biobank. Risk of these outcomes was compared between individuals orally taking a particular class of antihypertensive drug (in combination or as monotherapy) at recruitment with those not taking that type of medication. Analysis was restricted to White British individuals, including only incident events, exclusion of individuals with a diagnosis of the condition of interest prior to recruitment, and with adjustment made in the logistic regression analysis for age, sex, BMI, Townsend Deprivation Index, smoking status and a previous history of cardiovascular disease.

Statistical analysis was undertaken using R version 3.4.1 (The R Foundation for Statistical Computing) and Stata 14.2 (StataCorp LP). All supporting data are available within the article, its supplementary files and the web links provided. UK Biobank data was accessed through application 10035.

## Results

### Instrument selection

The genes, enhancer and promoter regions corresponding to the targets of each first-line antihypertensive drug are shown in Supplementary Table 1. We identified 1 gene for each drug target for ACEIs (*ACE*), ARBs (*AGTR1*), BBs (*ADRB1*) and TDs (*SLC12A3*), and 11 genes for CCBs (*CACNA1D, CACNA1F, CACNA2D1, CACNA2D2, CACNA1S, CACNB1, CACNB2, CACNB3, CACNB4*, *CACNG1, CACNA1C*), encoding the different calcium channel subunits. The *CACNA1F* gene is located on the X chromosome, and SNPs corresponding to this region were not available. Using the pre-defined instrument selection criteria, we identified 1 SNP for ACEIs, 6 SNPs for BBs and 24 SNPs for CCBs (Supplementary Tables 2-4). Therefore, as we identified 3 or more SNPs as instruments for BBs and CCBs, only these were taken forward to MR and PheWAS analyses. The F statistic for these SNPs ranged from 54 to 534 (Supplementary Tables 3-4), indicating a low risk of weak instrument bias (12).

### Mendelian randomisation

The main MR analysis using the 6 instruments for BBs identified a protective effect of genetically determined lower SBP through this target on CHD risk (RR 0.62, 95% confidence interval [CI] 0.47- 0.81, *P*=4×10^−4^). MR analysis using the same instruments did not identify an effect on stroke risk (RR 0.91, 95%CI 0.73-1.14, *P*=0.41). For CCBs, the main MR analysis using the 24 instrument SNPs identified a protective effect on CHD risk (RR 0.73, 95%CI 0.64-0.84, *P*=6×10^−6^) and stroke risk (RR 0.75, 95%CI 0.66-0.84, *P*=1×10^−6^). Individual MR estimates for each instrument SNP considered are given in Supplementary Figures 1-4. The MR estimates had overlapping 95% confidence intervals to those from RCTs of these drugs against placebo (Figure 1), considering a total of 19,313 participants for BBs and 4,695 participants for CCBs (3).

**Figure 1.**
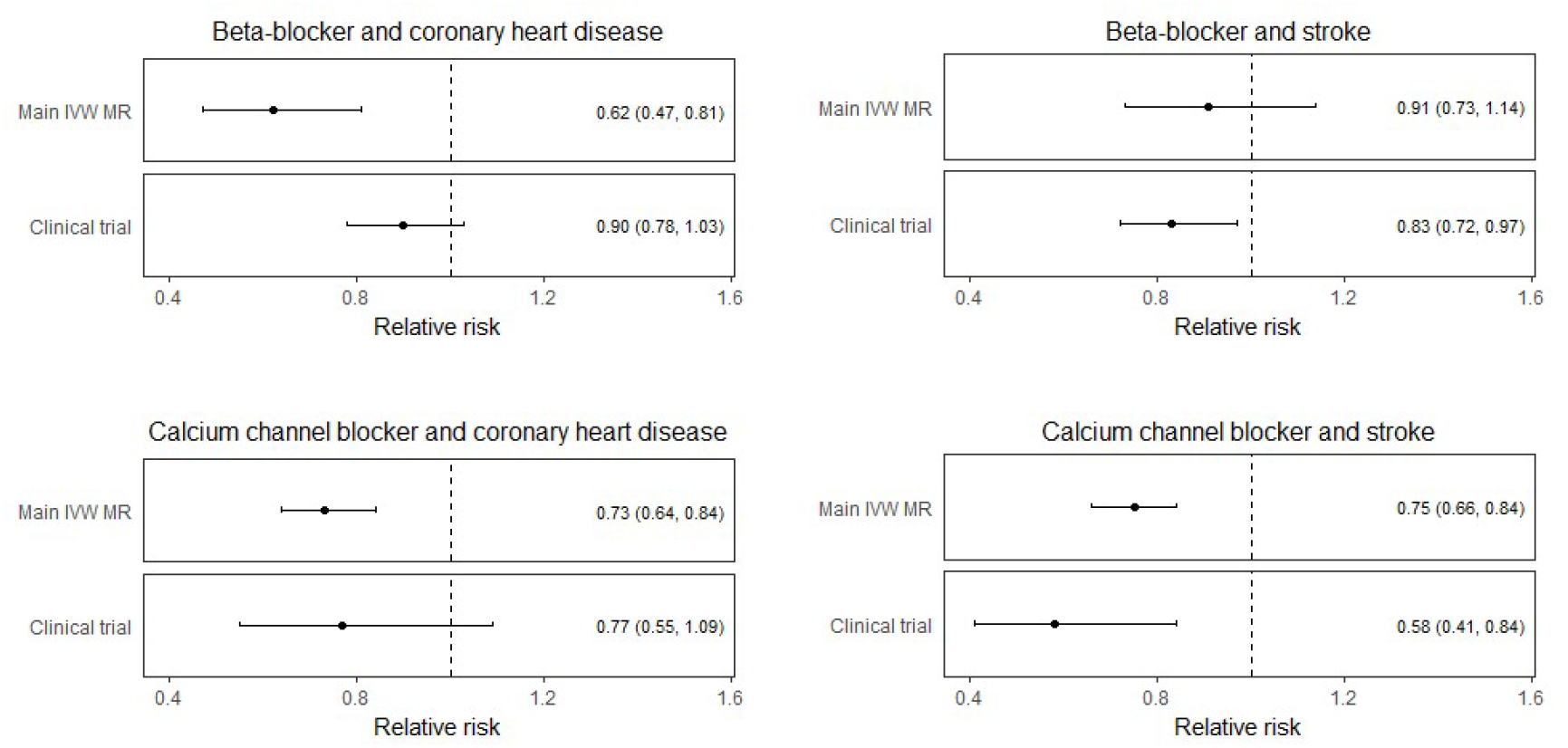
Mendelian Randomisation (MR) estimates for the effect of genetically lower systolic blood pressure through the beta-blocker and calcium channel blocker instruments respectively on risk of coronary heart disease and stroke, as compared to randomised controlled trial meta-analysis results (3). MR estimates are scaled to the effect of the corresponding drug in trial meta-analyses (3). IVW: inverse variance weighted.

The main instruments were based on genetic association estimates that corrected for antihypertensive medication use and adjusted for BMI (10). To avoid possible bias related to medication non-compliance or introduction of collider effects respectively, we performed sensitivity analyses using the UK Biobank SBP GWAS that did not correct for medication use or adjust for BMI (11). We identified 2 SNPs as instruments for BB (Supplementary Table 5), and 6 SNPs as instruments for CCB (Supplementary Table 6). IVW MR produced estimates that were comparable to the main analysis, but with wider confidence intervals (Supplementary Figures 5-8). Searching PhenoScanner (17, accessed on 6 August 2018), we identified possible pleiotropic effects through birthweight for the SNP rs1801253 used as an instrument for BBs, and through schizophrenia risk for the SNP rs714277 used as an instrument for CCBs. Repeating the IVW MR analysis after excluding these SNPs also produced similar estimates to the main analysis (Supplementary Figures 5-8).

We only found evidence of heterogeneity, suggesting possible bias related to pleiotropic SNPs, in the MR analysis of BBs on stroke risk (I^2^ 59%, Cochran’s Q *P*=0.03). The MR-Egger intercept was not significant for directional pleiotropy for either BBs (CHD *P*=0.87 and stroke *P*=0.89) or CCBs (CHD *P*=0.89 and stroke *P*=0.51). MR-PRESSO only detected outlier SNPs in the analysis of BBs on stroke risk (2 outliers), with MR-PRESSO estimates that excluded these SNPs consistent with the main analysis results (Supplementary Figure 6). Estimates using MR-Egger regression, the weighted median approach and MR-PRESSO also produced similar estimates to the main IVW MR analyses (Supplementary Figures 5-8).

### Phenome-wide association study

Following quality control and mapping of ICD-9 and ICD-10 to phecodes, data for 424,439 individuals across 909 distinct phenotypes were available for PheWAS analysis. Using the main instruments, the PheWAS for BBs revealed associations with a lower risk of hypertension, circulatory disease and atrial fibrillation and flutter (Figure 2, Supplementary Table 7). The GRS for CCBs was also associated with a lower risk of traits related to hypertension and cardiovascular disease (Figure 4, Supplementary Table 8). CCBs additionally showed an association with higher risk of diverticulosis (OR 1.23, 95%CI 1.10-1.38, *P*=2.39×10^−4^). In sensitivity analyses using instruments derived from the SBP GWAS that did not correct for medication use or adjust for BMI (Supplementary Tables 9-10), or after excluding the potentially pleiotropic rs1801253 and rs714277 SNPs (Supplementary Tables 11- 12), similar estimates were obtained to the main analysis.

**Figure 2.**
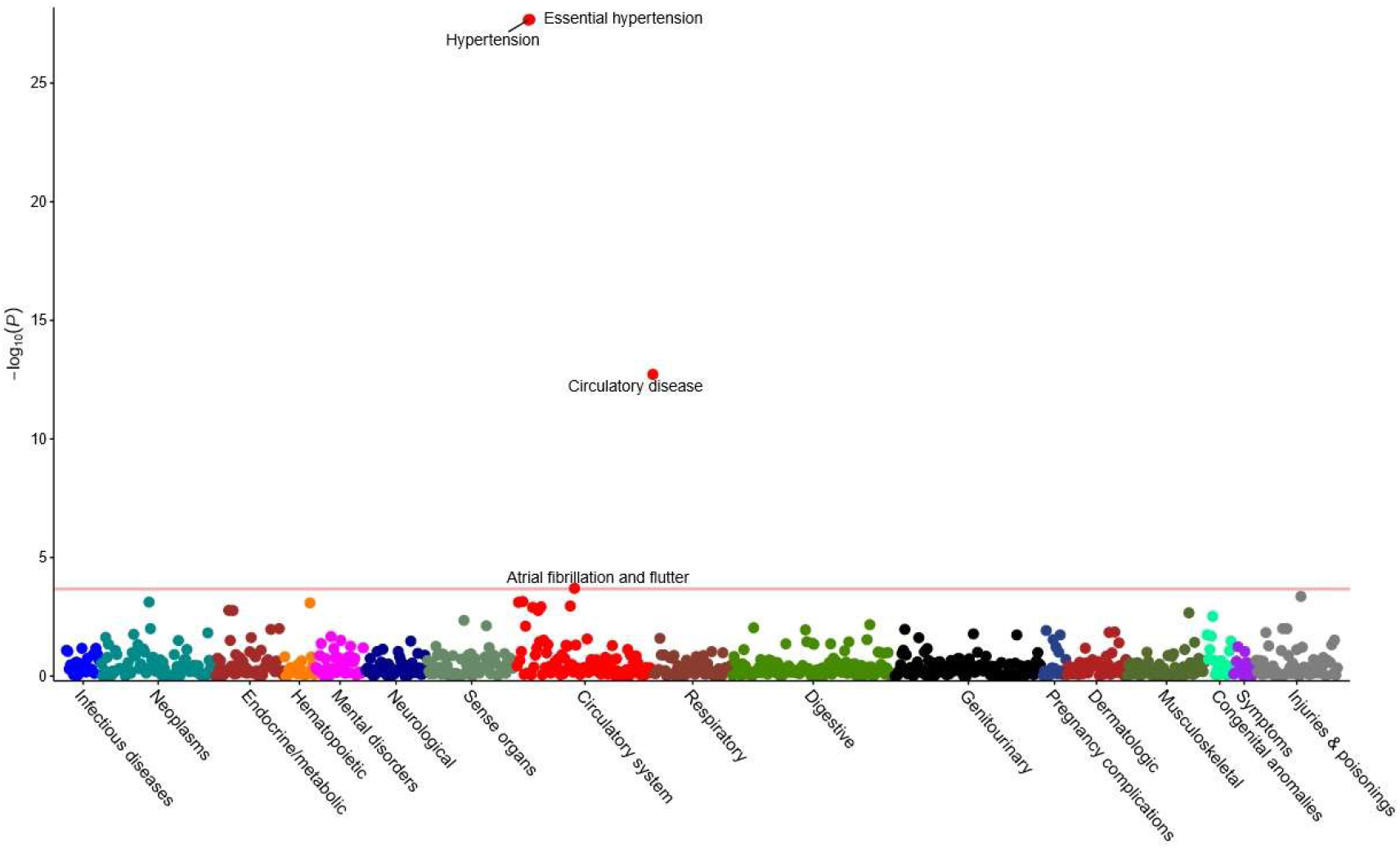
Phenome-wide association study (PheWAS) in the UK Biobank of the genetic risk score for beta-blockers. Phenotypes significant at a false discovery rate threshold of 5% are detailed.

**Figure 3.**
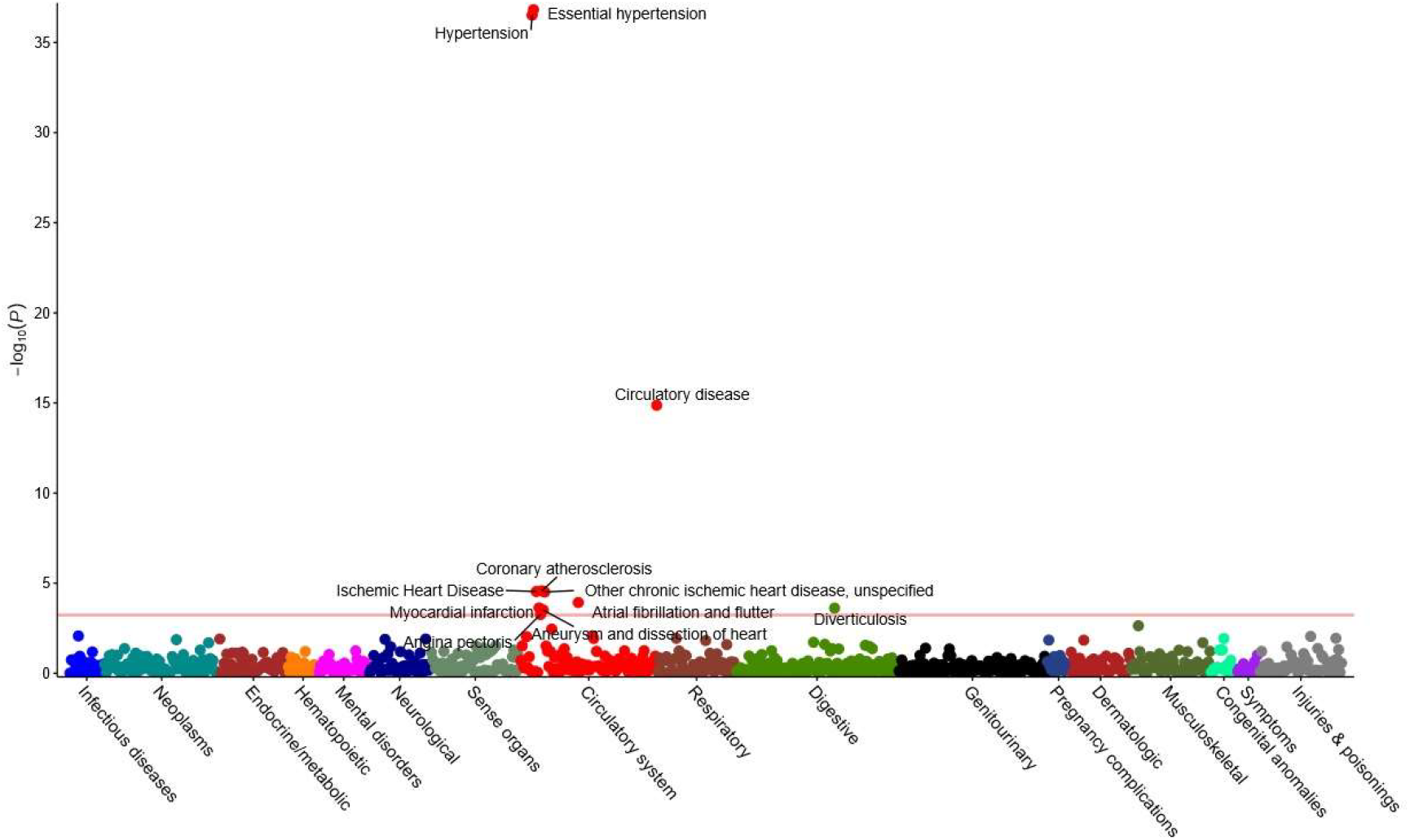
Phenome-wide association study (PheWAS) in the UK Biobank of the genetic risk score for calcium channel blockers. Phenotypes significant at a false discovery rate threshold of 5% are detailed.

**Figure 4.**
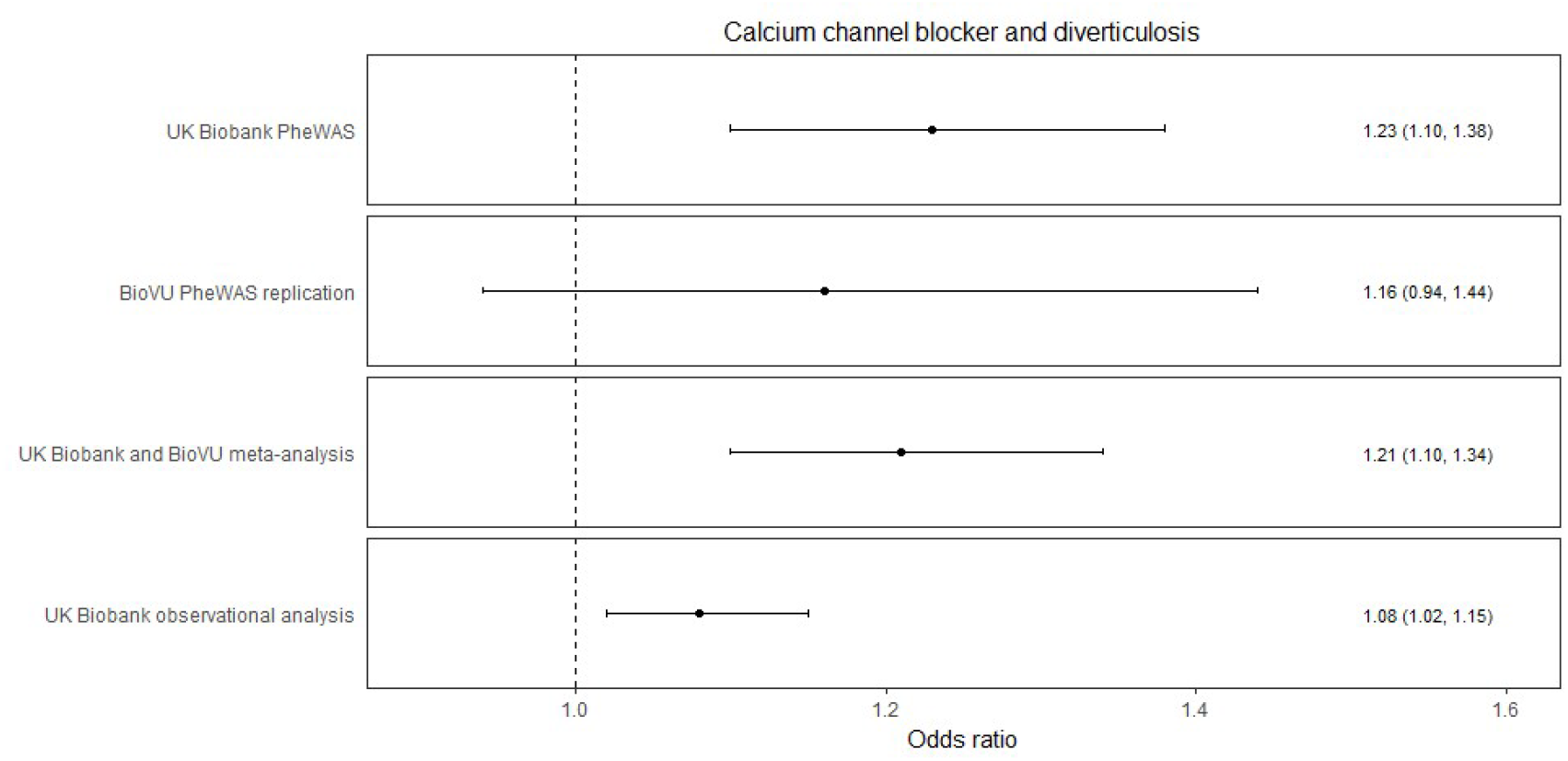
A forest plot comparing estimates for the association between calcium channel blockers and diverticulosis risk derived from phenome-wide association study (PheWAS) analyses in the UK Biobank and Vanderbilt University Biobank (BioVU) respectively, their fixed-effects pooled estimate, and observational analysis in the UK Biobank. PheWAS results are scaled to the SBP lowering effect of CCBs in clinical trials (3), while observational analysis refers to the effect of CCB use.

Data for 45,517 individuals were available in BioVU to further investigate novel PheWAS findings for traits unrelated to hypertension. The prevalence of diverticulosis in BioVU was 12%, comparable to the 10% observed in the UK Biobank. In BioVU, the CCB GRS association with diverticulosis had OR 1.16 (95%CI 0.94-1.44, *P*=0.17), and the meta-analyses of UK Biobank and BioVU strengthened the overall evidence for an association (OR 1.21, 95%CI 1.10-1.34, *P*=1.14×10^−4^). Although the confidence interval for the association in BioVU alone crossed the null, the effect estimate was similar to that observed in the UK Biobank PheWAS (Supplementary Table 13 and Figure 4).

### Observational analysis

For the observational analysis of diverticulosis in the UK Biobank, there were 495,676 individuals left after excluding those with a diagnosis prior to recruitment, with 10,955 incident cases and 35,626 individuals taking CCBs. After adjusting for confounders, there was a significant association between CCB use and risk of diverticulosis (OR 1.08, 95%CI 1.02-1.15, *P*=0.01), consistent with that identified in the PheWAS analysis (Figure 4).

## Discussion

We leveraged large-scale GWAS data from over 750,000 individuals and generated genetic instruments for the effect of BBs and CCBs, two of the most commonly used medications worldwide. The MR estimates for risk of CHD and stroke were comparable to those observed in RCTs for BBs and CCBs respectively against placebo, supporting the validity of our instruments. Notably, MR also found CCBs to have a protective effect on risk of stroke but did not provide evidence of this for BBs (Figure 1), in keeping with the increased blood pressure variability observed with BBs and their lower effectiveness for preventing stroke (26). PheWAS on 909 outcomes corroborated the known efficacy of these agents in preventing hypertension and related vascular diseases, thus further supporting the robustness of the instruments used.

The PheWAS investigation also revealed an increased risk of diverticulosis associated with the GRS for CCBs. A similar estimate was found in BioVU, which contained fewer cases and had a correspondingly wider confidence interval that crossed the null. Meta-analysis of estimates from the UK Biobank and BioVU further strengthened the significance of the association (Figure 4). Our finding was supported by observational analysis showing that CCB treatment at baseline in UK Biobank was associated with increased risk of diverticulosis. In terms of a possible mechanism, constipation is an established side effect of CCBs related to their role in reducing bowel contractility (27), and it maybe through a similar process that the risk of diverticulosis is increased. The same phenomenon may also paradoxically lower the risk of bowel perforation within the context of diverticular disease (27). Diverticulosis is one of the most common reasons for hospital admission in the United States (28), and is rising in incidence (29). With over a tenth of the world’s adults estimated to have hypertension (1), and the recommendation of CCBs as a first-line pharmacological agent (6), the clinical implications of these findings merit consideration. For example, individuals with diverticulosis or at increased risk of developing it may benefit from alternative pharmacological treatments for hypertension.

The genetic instruments for BBs and CCBs did not show detrimental associations with any of the other traits examined in PheWAS. While absence of evidence is not evidence of absence, this does provide some assurance that long-term pharmacological inhibition of these drug targets is generally safe, with other side effects that require hospitalisation being smaller or rarer.

A major strength of our work is that it uses genetic variants to instrument the effect of antihypertensive drugs using existing data obtained from large-scale studies, thus avoiding the time and resource constraints associated with such study through RCTs (4), and additionally overcoming the limitations of potential confounding and reverse causation from use of standard observational methods (7). We performed a range of sensitivity analyses to support the robustness of our findings, including investigation of possible ascertainment bias relating to medication non-compliance or collider bias arising from the BMI adjustment in the genetic association estimates for SBP (10). Similar consistency with our main findings was seen in the sensitivity analyses that excluded potentially pleiotropic SNPs or when applying statistical methods that are more robust to the inclusion of pleiotropic variants. Additionally, PheWAS allowed rapid investigation of hundreds of traits across the phenome, with opportunity for further investigation in an independent cohort, as well as in observational analysis.

Concerning limitations, the MR and PheWAS results estimate the cumulative effect of lifelong exposure to genetic variants, rather than the consequence of a clinical intervention. This is in keeping with our PheWAS identifying a larger effect of CCBs on risk of diverticulosis than the observational analysis (Figure 4). Furthermore, there may be unknown pleiotropic effects of the genetic variants that bias the association estimates (13). Sample overlap between the populations used to derive instruments and measure genetic association estimates for outcomes can also introduce bias when weak instruments are used (30), although this did not appear to be a feature of our data. While less stringent criteria for selecting instruments (such as a more relaxed *P*-value threshold for association with SBP, or a more lenient LD criterion for clumping) may have increased the number of variants available, this may also have reduced the sensitivity and specificity of the analysis due to introduction of weak instrument bias and invalid instruments respectively. Similarly, taking targets for which there were fewer than 3 instrument SNPs forward to MR and PheWAS analyses would not have allowed for statistical evaluation of possible bias related to pleiotropy (13), and could have thus risked generating misleading results.

In conclusion, we have identified genetic variants that are able to instrument the effect of the BB and CCB classes of antihypertensive medication. In MR and PheWAS, our instruments corroborated the established associations of these agents with a range of traits related to hypertension. Additionally, we identified an apparent, previously unreported detrimental effect of CCBs on risk of diverticulosis that was supported in observational analysis, a finding that requires further replication before it should alter clinical practice. We did not identify any other potential side effects of either drug class to suggest a lack of long-term safety. Our study demonstrates that the use of genetic variants offers a powerful complement to existing RCT and observational approaches for investigating the efficacy, side effects and repurposing potential of antihypertensive agents.

## Contributors

DG, IT, MKG and MD designed the study. DG, MKG, FK and LJ performed the analysis. All authors interpreted the results. DG and IT drafted the manuscript. All authors critically revised the manuscript for intellectual content. All authors approved the submitted version and are accountable for the integrity of the work.

## Acknowledgements

DG is funded by the Wellcome 4i Clinical PhD Programme at Imperial College London. MG is funded by scholarships from the Deutscher Akademischer Austauschdienst and the Onassis Foundation. QF, W-QW and JCD are funded by National Institute of Health grants R01 HL133786, R01 GM120523, R01 LM010685. ET acknowledges support from Cancer Research UK (C31250/A22804). PE acknowledges support from the Medical Research Council and Public Health England (MR/L01341X/1) for the Medical Research Council-Public Health England Centre for Environment and Health; the National Institute for Health Research Imperial Biomedical Research Centre in collaboration with Imperial College NHS Healthcare Trust. PE is supported by the UK Dementia Research Institute, which receives funding from UK DRI Ltd funded by the UK Medical Research Council, Alzheimer’s Society and Alzheimer’s Research UK. PE is associate director of the Health Data Research UK London funded by a consortium led by the UK Medical Research Council. MD acknowledges funding from the European Union’s Horizon 2020 research and innovation programme under grant agreements No 666881, SVDs@target and No 667375, CoSTREAM; the DFG as part of the Munich Cluster for Systems Neurology (EXC 1010 SyNergy) and the CRC 1123 (B3); the Corona Foundation; the Fondation Leducq (Transatlantic Network of Excellence on the Pathogenesis of Small Vessel Disease of the Brain); the e:Med program (e:AtheroSysMed) and the FP7/2007-2103 European Union project CVgenes@target (grant agreement number Health-F2-2013-601456). The MEGASTROKE project received funding from sources specified at http://www.megastroke.org/acknowledgments.html. Details of all MEGASTROKE authors are available at http://www.megastroke.org/authors.html.

The authors acknowledge the contributors of the data used in this work: CARDIoGRAMplusC4D, International Consortium for Blood Pressure, MEGASTROKE, UK Biobank and Vanderbilt University Biobank. Links to the various web resources used are provided below.

CARDIoGRAMplusC4D: http://www.cardiogramplusc4d.org/

DrugBank: https://www.drugbank.ca/

GeneCards: https://www.genecards.org/

MEGATSROKE GWAS meta-analysis summary data: http://www.megastroke.org/

Neale Labe UK Biobank GWAS summary data: http://www.nealelab.is/blog/2017/7/19/rapid-gwas-of-thousands-of-phenotypes-for-337000-samples-in-the-uk-biobank

PhenoScanner: www.phenoscanner.medschl.cam.ac.uk/phenoscanner

UK Biobank: http://www.ukbiobank.ac.uk/

Vanderbilt University Biobank: https://victr.vanderbilt.edu/pub/biovu/

